# Meta-analysis reveals taxon- and life stage-dependent effects of ocean acidification on marine calcifier feeding performance

**DOI:** 10.1101/066076

**Authors:** Jeff C. Clements

## Abstract

While ocean acidification is considered among the greatest threats to marine ecosystems, its effects on the feeding performance of marine calcifiers remain uncertain. I conducted a meta-analysis of effect sizes (LnRR) assessing the impacts of acidification on the feeding ability of three groups of marine calcifiers - molluscs, arthropods, and echinoderms. Results suggested taxon-dependent effects of acidification on calcifier feeding performance, with depressed feeding observed for molluscs, echinoderms, and when all taxa were considered. However, ocean acidification had no effect on feeding performance in marine arthropods and larval feeding performance appeared more vulnerable than that of juveniles and adults. Feeding performance was not related to acclimation time nor *p*CO_2_ level. This study suggests that the feeding performance of molluscs and early life-stage echinoderms may be depressed in a more acidic ocean, but that arthropod feeding performance is unlikely to suffer. Such changes in feeding performance could contribute to slower growth and development in the early life stages of these organisms and could potentially contribute to changes in community and ecosystem structure where these organisms coexist. Finally, feeding performance could, at least in part, moderate the degree to which molluscs and echinoderms can use food to overcome acidification effects early in life.

## Introduction

Since the Industrial Revolution, anthropogenic activity, primarily the burning of fossil fuels, has contributed to an unprecedented increase in atmospheric carbon dioxide (CO_2_) concentrations [1-3]. While terrestrial consequences are evident (i.e., global warming), the anthropogenic increase in atmospheric CO_2_ concentrations has predictable outcomes on physicochemical processes in the marine environment [4]. One such outcome of elevated atmospheric CO_2_ is a process known as ocean acidification. In brief, ocean acidification refers to the reduction of oceanic pH as a result of absorbing excess anthropogenic atmospheric CO_2_. The absorption of excess atmospheric CO_2_ by oceanic surface waters has reduced surface-ocean pH by 0.1 units since the Industrial Revolution, with an expected further 0.2-0.3 unit drop by the end of the century [1].

Ocean acidification is suspected to be among the greatest threats to marine ecosystems globally [4-5]. In particular, acidification has been reported to impact physiological functioning [6-7] and animal behaviour [8-10] in a wide variety of marine organisms in both positive and negative ways, potentially leading to altered population-, community-, and ecosystem functioning. Although considered a major threat to marine systems and receiving widespread attention over the past decade, the large-scale effects of ocean acidification on many biological processes remain poorly understood.

Although ocean acidification is likely to impact marine calcifiers in the future [7], it has recently been suggested that these organisms may be able to overcome physiological stress associated with acidification given adequate food supply [11-13]. Experimental studies exploring this hypothesis typically rear animals under varying CO_2_ conditions and food availabilities over some period of time, measuring biological endpoints (e.g., calcification and growth) before and after rearing. However, such studies have thus far neglected to incorporate the impacts of acidification on post-rearing feeding performance and other aspects of energy acquisition into the equation, many of which are known to be affected by ocean acidification [14-17]. Feeding performance is critical to an animal’s ability to acquire energy and is likely to moderate, at least in part, the degree to which calcifiers will be able to overcome acidification stress. While recent meta-analyses suggest a moderating effect of food availability on calcification and growth in marine calcifiers [13], taxonomic effects were not tested (presumably due to a limited dataset). It is thus important to understand the ways in which the feeding performance of these organisms will be impacted under future oceanic conditions across calcifying taxa in order to accurately predict the biological consequences of ocean climate change.

Given that numerous studies assessing the impact of ocean acidification on the feeding performance of marine organisms have been accumulated over the past decade, a synthesis of such impacts is warranted. Here I present the results of a meta-analysis aimed at determining the impacts of ocean acidification on the feeding performance of three groups of marine calcifiers – molluscs, arthropods, and echinoderms. Results are discussed in the context of community- and ecosystem-wide impacts and in relation to the ability of calcifiers to overcome acidification stress by utilizing food availability.

## Methods

Articles were searched for via Web of Science and Google Scholar using the keywords “ocean acidification” *plus* feeding *and/or* ingestion *and/or* consumption (anywhere in article). I then conducted a cited literature search of each article obtained from the online literature search to collect any articles that may have been missed in the online search. I considered feeding/grazing rates, ingestion rates, absorption rates, and food mass loss as appropriate measures of feeding. The highest pH level (or lowest *p*CO_2_ level) employed in each experiment from each study was considered the control treatment, and I only considered acidification conditions relevant to near-future predictions (Table 1). For studies reporting different acclimation periods or experimental durations, I used only the longest period/duration since future conditions will be long-lasting. If studies used > 1 experimental assay, species, size, relevant CO_2_ concentration, or food supply, I treated each as an individual observation. For experiments that tested the combined effects of acidification and other environmental stressors (i.e., warming and hypoxia) on feeding performance, I only assessed differences between acidification groups for the control levels of the other factors (i.e., ambient temperature and O_2_; based on authors’ definition provided in each study).

**Table 1.** General information collected from each of the 21 studies used in the meta-analysis of ocean acidification effects on marine calcifier feeding performance. Bolded references indicate studies that assessed the combined effects of ocean acidification and warming. Studies are ordered chronologically within their respective Phylum.

| Phylum | Reference | Acclimation time (d) | Species | Life stage | Geographic location | pCO<sub>2</sub> | pH | Outcome |
| --- | --- | --- | --- | --- | --- | --- | --- | --- |
| Mollusca | [26] | 70 | *Mytilus chilensis* | Juvenile | Chiloé, southern Chile | 750 ppm | 7.71 | - |
|  |  | 70 |  |  |  | 1200 ppm | 7.57 | - |
|  | [27] | 78 | *Mytilus galloprovincialis* | Juvenile | Ría de Ares-Betanzos, Spain | 1989 µatm | 7.74 | = |
|  | [28] | **150** | ***Littorina littorea*** | **Adult** | **Plymouth, England** | **750 µatm** | **-** | **=** |
|  |  | **150** |  |  |  | **1000 µatm** | **-** | **=** |
|  | [29] | **42-56** | *Concholepas concholepas* | Larvae | Las Cruces, Chile | **700 ppm** | **7.67** | - |
|  |  | **42-56** |  |  |  | **1000 ppm** | **7.52** | - |
|  | [30] | **70** | *Concholepas concholepas* | Larvae | Las Cruces, Chile | **700 ppm** | **7.80** | - |
|  |  | **70** |  |  |  | **1000 ppm** | **7.65** | - |
|  |  | **70** | *Perumytilus purpuratus* | Juvenile | Las Cruces, Chile | **700ppm** | **7.87** | - |
|  |  | **70** |  |  |  | **1000 ppm** | **7.73** | - |
|  | [17] | 3 | *Mytilus coruscus* | Juvenile | Zhejiang Province, China | 1023 uatm | 7.7 | = |
| Arthropoda | [30] | 210 | *Palaemon pacificus* | Adult | Nagasaki, Japan | 1000 ppm | 7.89 | = |
|  | [31] | 70 | *Carcinus maenas* | Adult | Kiel Fjord, Baltic Sea | 1247 µatm | 7.84 | = |
|  | [32] | 1 | *Euphausia superba* | Adult | Adelaide Island, Antarctica | 672 ppm | 7.84 | + |
|  | [33] | **27** | ***Pandalus borealis*** | **Larvae** | **Hillefjord, Norway** | **1038-1751 µatm** | **7.6-7.65** | **=** |
|  | [34] | **14** | ***Peramphithoe parmerong*** | **Adult** | **New South Wales, Australia** | **-** | **7.60** | **=** |
|  | [35] | 74 | *Calanus finmarchicus* | Adult | Trondheim, Norway | 1080 µatm | 7.65 | = |
|  | [36] | 71 | *Panopeus herbstii* | Adult | Beaufort, North Carolina | 785 µatm | 8.04 | - |
|  | [37] | **1** | ***Tigriopus japonicus*** | **Adult** | **Xiamen Bay, China** | **1007 µatm** | **7.80** | **=** |
|  | [38] | 10 | *Pseudocalanus acuspes* | Adult | Skagerrak, Norway | 823-853 µatm | 7.70-7.80 | = |
|  |  | 10 |  |  |  | 1315-1468 µatm | 7.47-7.61 | = |
| Echinodermata | [39] | **14** | ***Pisaster ochraceus*** | **Juvenile** | **Vancouver, British Columbia** | **780 ppm** | **-** | **+** |
|  | [40] | 20 | *Strongylocentrotus purpuratus* | Larvae | Pasadena, California | 1271 µatm | 7.70 | - |
|  | [31] | 70 | *Asterias rubens* | Adult | Kiel Fjord, Germany | 1247 µatm | 7.84 | = |
|  | [41] | **6** | ***Amblypneustes pallidus*** | **Adult** | **Lady Bay, South Australia** | **640-650 µatm** | **7.93** | **+** |
|  | [42] | 14 | *Asterias rubens* | Juvenile | Antigonish, Nova Scotia | - | 7.94 | - |
|  | [16] | 1200 | *Sterechinus neumayeri* | Adult | Ryder Bay, Antarctica | 834 µatm | 7.72 | = |
|  |  | 1200 |  |  |  | 1361 µatm | 7.52 | = |

From each study, I collected general information for each species tested, including their associated Phylum, their life history stage (larvae, juvenile, or adult), and the duration that they were reared under experimental or control conditions prior to the onset of the experiment (hereafter referred to as “acclimation time”, measured in days). I also collected general study information, including the geographic location from which the animals were collected and the pH and/or *p*CO_2_ conditions of each relevant treatment. The overall outcome of ocean acidification on feeding performance for each study was also recorded. This was determined by the statistical outcome of each study, whereby α ≤ 0.05 was used as a threshold to define elevated (acidification feeding performance > control) and depressed (acidification feeding performance < control) feeding performance (or overall outcome); α > 0.05 defined no change in feeding performance. For studies that employed multiple experiments with different statistical outcomes, the statistical outcome most often observed was deemed as the overall outcome for that study. In order to determine individual and mean effect sizes, I recorded the mean, variance (standardized as standard deviation), and sample size in feeding performance for each relevant acidification treatment and the associated control treatment for each observation in each study. These values were obtained from published tables, were digitally estimated from published graphs using ImageJ, or were obtained from online databases. Means, variances, and sample sizes were used to calculate natural log response ratios (effect size, LnRR) for each relevant experimental group. LnRR is the ratio of the experimental effect to the control effect (natural log transformed). An LnRR value of 0 signifies no effect of the experimental treatment on the response variable, while negative and positive LnRR values signify negative and positive effects, respectively [18]. I used LnRR as an effect size because this metric has a high capacity to detect true effects and is robust to low sample sizes [19].

Weighted random effects models were used to derive mean effects sizes (LnRR) for each experimental treatment and for each taxonomic group and life history stage (see Results); mean effect sizes were only calculated for taxa and/or life history stages containing 3 or more observations (n ≥ 3). Bootstrapped (10,000 replicates) bias corrected and accelerated (BCa) 95% confidence intervals were used to determine statistical significance (significance occurs when 95% confidence interval does not cross 0). Q tests (α ≤ 0.05) were used to test for heterogeneity in effect sizes among studies. Linear regression was used to determine if *p*CO_2_ levels and/or acclimation time affected feeding performance (as measured by LnRR). All analyses were conducted using R v. 3.2.4 [20] (see Table S1 for R code); effect size calculations and analyses were conducted using the Metafor package [21].

## Results

### Literature search

The literature search resulted in a total of 21 studies, encompassing 19 species spanning 3 taxonomic groups: 6 species of molluscs, 10 species of arthropods, and 5 species of echinoderms (Table 1). A total of 52 overall observations (i.e., data points) were used in the analyses, including 17 mollusc observations, 25 arthropod observations, and 10 echinoderm observations.

### Overall study outcomes

In general, studies reported mixed effects of ocean acidification on marine calcifier feeding performance (Figure 1). When all taxa were considered, ∼30% of studies reported depressed feeding performance under ocean acidification, while ∼15% reported enhanced feeding performance; ∼55% of studies reported no change in feeding performance. Similarly, 33% of echinoderm studies reported depressed feeding performance under ocean acidification, while another 66% reported either enhanced feeding (33%) or no change (33%). For molluscs, 50% of studies reported depressed feeding performance under ocean acidification, while the other 50% found no change. Arthropods exhibited enhanced feeding under ocean acidification in 11% of studies and depressed feeding in another 11%, while 78% of arthropod studies found no change in feeding performance under ocean acidification.

**Fig. 1.**
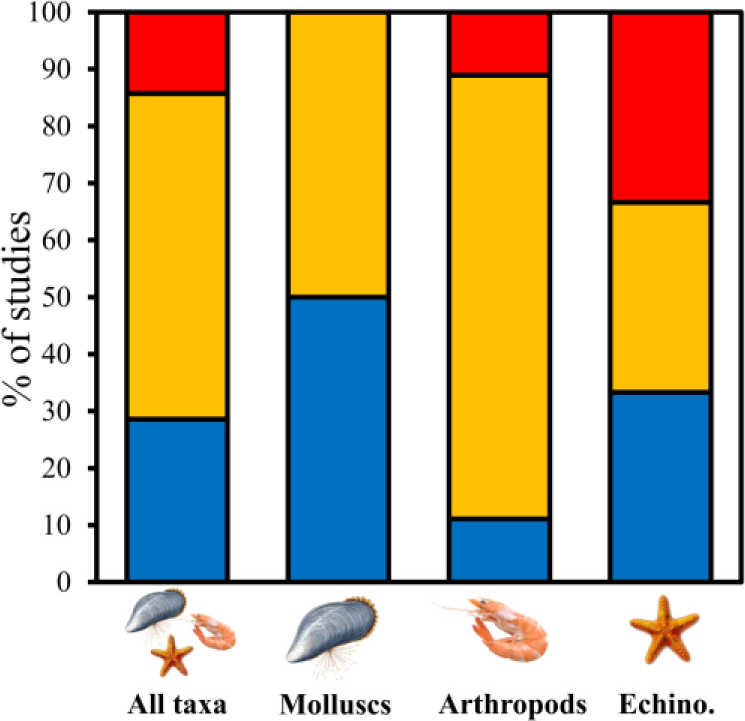
Percentage of overall outcomes of studies (n = 21) assessing the impacts of ocean acidification on marine calcifier feeding performance, assembled by taxa. Outcomes were either enhanced feeding (red bars; significantly lower feeding under acidification than in control), depressed feeding (blue bars; significantly higher feeding under acidification than in control), or no change (yellow; no significant difference between acidification and control).

### Effect size analysis

Effect sizes were significantly heterogeneous for each taxonomic grouping and life history stage, except for larval arthropods, and adult molluscs and echinoderms (Table 2). Considering all experimental observations from all studies, effect sizes appeared predominantly negative (Figure 2). Effect sizes of molluscan feeding performances were unanimously negative, with 100% of observations eliciting negative effect sizes. In contrast, arthropods and echinoderms each showed a ∼3:2 ratio of negative to positive effect sizes. When all taxa were considered, ∼70% and 30% of observations resulted in negative and positive effect sizes, respectively. Effect sizes were not correlated with acclimation time nor *p*CO_2_ level (Figure 3, Table 3).

**Fig. 2.**
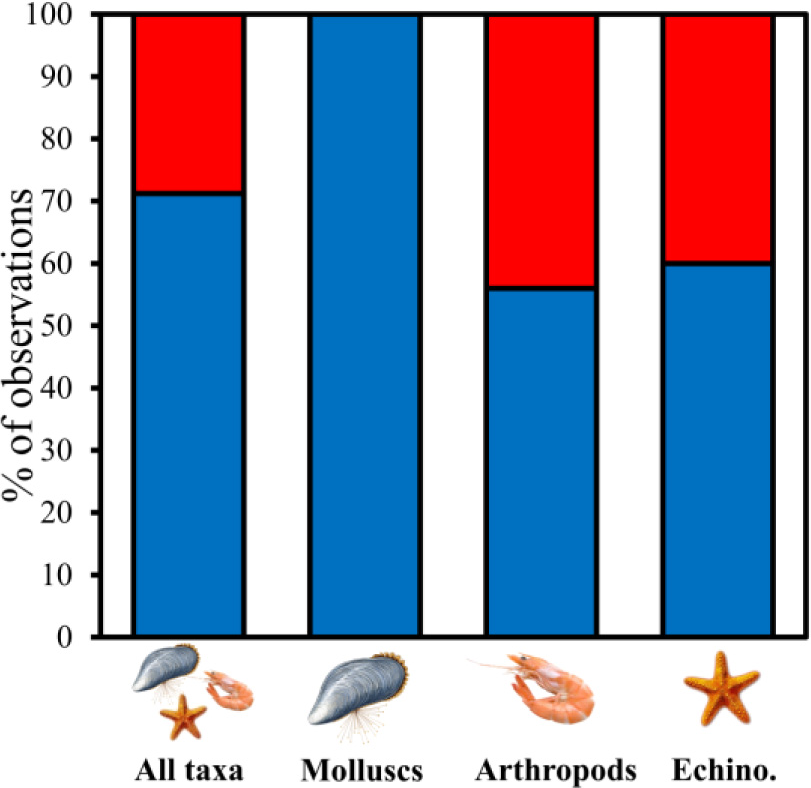
Percentage of total observations (n = 52) that elicited positive (red bars; LnRR > 0) and negative (blue bars; LnRR < 0) effect sizes.

**Table 2.** Results of Q-tests for heterogeneity of effect sizes.

| Phylum | All stages | Larvae | Juveniles | Adults |
| --- | --- | --- | --- | --- |
| All | $Q_{51} = 1159.8, P < 0.001$ | $Q_9 = 602.4, P < 0.001$ | $Q_8 = 104.5, P < 0.001$ | $Q_{32} = 151.1, P < 0.001$ |
| Mollusca | $Q_{16} = 904.2, P < 0.001$ | $Q_5 = 530.9, P < 0.001$ | $Q_6 = 24.3, P < 0.001$ | $Q_3 = 4.0, P = 0.263$ |
| Arthropoda | $Q_{24} = 111.0, P < 0.001$ | $Q_2 = 0.6, P = 0.732$ | - | $Q_{21} = 102.8, P < 0.001$ |
| Echinodermata | $Q_9 = 78.8, P < 0.001$ | - | - | $Q_6 = 4.3, P = 0.631$ |

**Fig. 3.**
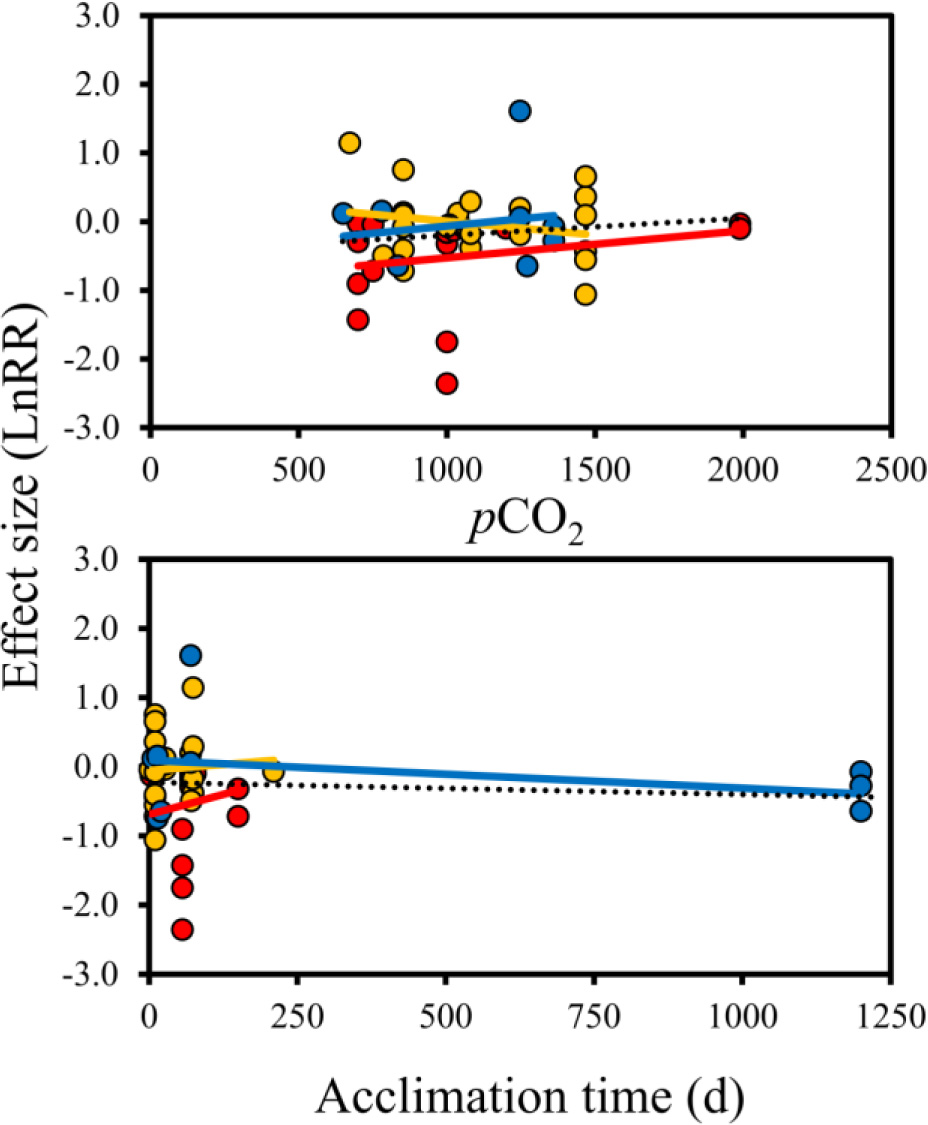
Linear relationships of effect sizes as a function of *p*CO_2_ level (top) and acclimation time (in days; bottom). Each point represents the effect size (LnRR) an individual observation for molluscs (red plots; n = 17), arthropods (yellow plots; acc. time n = 25, *p*CO_2_ n = 24), and echinoderms (blue plots; acc. time n = 10, *p*CO_2_ n = 9). Dashed black lines represent the linear relationship of all taxa pooled. See Table 3 for linear regression results.

**Table 3.** Results of linear regressions for relationships of calcifier feeding performance (effect sizes, LnRR) as a function of acclimation time (days) and *p*CO_2_ level.

| Predictor | Phylum | Equation | $R^2$ | df | F | P-value |
| --- | --- | --- | --- | --- | --- | --- |
| Acclimation time | All | $y = -0.0002x - 0.18$ | 0.01 | 51 | 0.49 | 0.49 |
| | Mollusca | $y = 0.0023x - 0.69$ | 0.01 | 16 | 0.19 | 0.66 |
| | Arthropoda | $y = 0.0007x - 0.06$ | $< 0.01$ | 24 | 0.11 | 0.74 |
| | Echinodermata | $y = -0.0004x + 0.09$ | 0.12 | 9 | 1.11 | 0.32 |
| $p\text{CO}_2$ | All | $y = 0.0002x - 0.47$ | 0.01 | 49 | 0.59 | 0.45 |
| | Mollusca | $y = 0.0004x - 0.92$ | 0.05 | 16 | 0.82 | 0.38 |
| | Arthropoda | $y = 0.0004x + 0.41$ | 0.05 | 23 | 1.07 | 0.31 |
| | Echinodermata | $y = 0.0004x - 0.48$ | 0.03 | 8 | 0.21 | 0.66 |

When all taxa were considered, marine calcifier feeding performance was significantly depressed by ocean acidification (Figure 4). When separated into respective taxonomic groups, however, only molluscs and echinoderms exhibited depressed feeding performance, while arthropod feeding was unaffected (Figure 4).

**Fig. 4.**
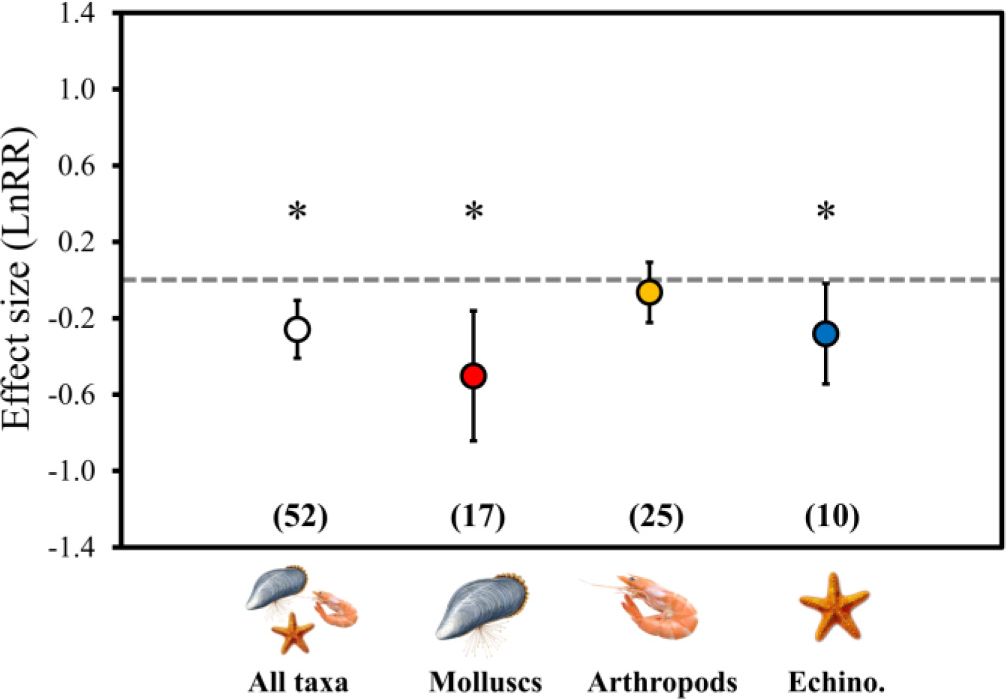
Mean effect size (LnRR) ± 95% confidence intervals (CI) for the effects of ocean acidification on feeding performance in the three taxonomic groups assessed in this study. Data were obtained from 21 published studies of 20 species assessing the effects of acidification on calcifier feeding (see Table 1). Asterisks indicate statistically significant effects. Numbers in parentheses indicate the number of observations used to generate the mean effect size and 95% CI.

When all taxa were considered, larval and juvenile calcifiers had depressed feeding performances under ocean acidification, although larvae elicited the most dramatic response; adults were unaffected (Figure 5). In molluscs, all life stages showed depressed feeding performance under ocean acidification, while larval and adult arthropods, along with adult echinoderms, were unaffected. I was unable to test for effects on juvenile arthropods and larval and juvenile echinoderms, due to an insufficient number of observations (< 3).

**Fig. 5.**
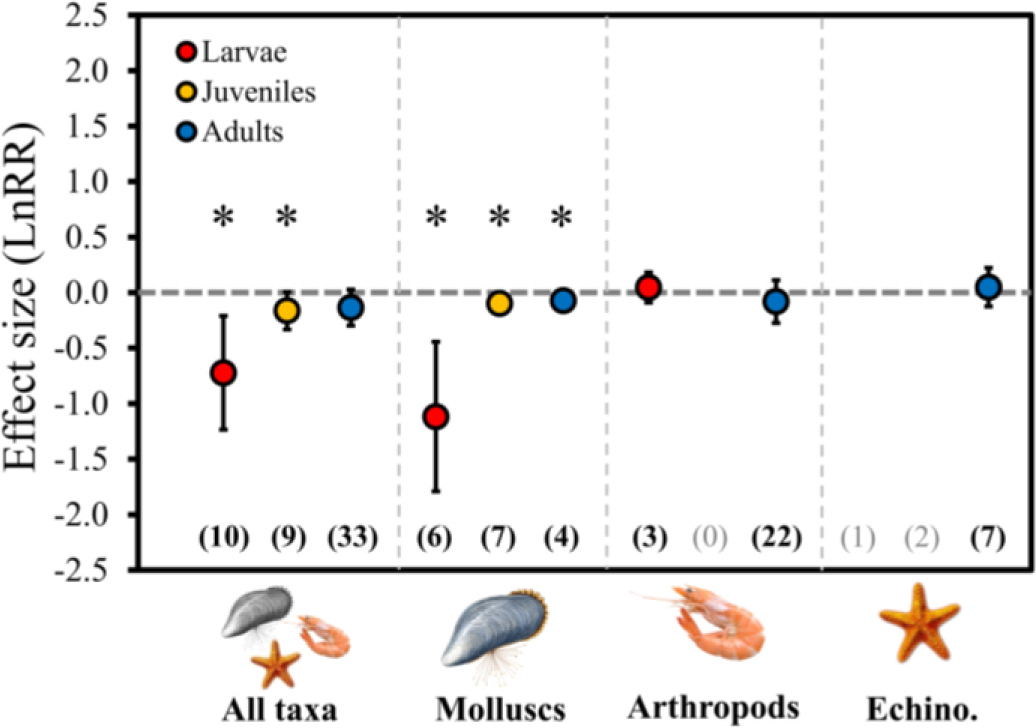
Mean effect size (LnRR) ± 95% confidence intervals (CI) for the effects of ocean acidification on feeding performance in the three taxonomic groups assessed in this study.. Data were obtained from 21 published studies of 20 species assessing the effects of acidification on calcifier feeding (see Table 1). Asterisks indicate statistically significant effects. Numbers in parentheses indicate the number of observations used to generate the mean effect size and 95% CI.

## Discussion

This study suggests that ocean acidification has the capacity to impact the feeding performance of marine calcifiers, although effects are dependent upon taxa and life-stage (but not *p*CO_2_ level or duration of acclimation). When grouped together by life stage and taxa, marine calcifiers exhibited lower feeding performance under near-future (end of century) ocean acidification conditions. However, when broken down by taxa, it appeared that while molluscs and echinoderms experience poorer feeding performance under higher CO_2_ conditions, arthropods are unaffected. In addition, when further broken down by life-history stage, larval feeding performance was much more susceptible to the effects of acidification than juveniles and adults, although juvenile and adult molluscs were still impacted. Given the contrasting results between taxa and life history stage, it is plausible that ocean acidification effects to feeding performance could contribute to changes in the overall structure and function on marine invertebrate communities. For example, in a more acidified ocean, depressed feeding by filter feeding bivalves could result in increased plankton biomass and contribute to increased prevalence of plankton blooms and potential hypoxic events. In addition, depressed feeding in molluscs could lead to poorer-quality molluscan prey for predatory crabs, resulting in crabs needing to eat more to maintain an adequate energy budget. Given that crab feeding performance appears unaltered in a more acidified ocean, one might predict that current densities of mussels residing in the presence of predatory arthropods may decrease. Such scenarios become more complicated in systems where various predators spanning multiple taxa (i.e., molluscs, arthropods, and echinoderms) are consuming a similar prey source like filter feeding bivalves. Furthermore, additional effects of ocean acidification (e.g., behavioural impacts to foraging and prey defenses, other physiological responses) will almost certainly contribute to predatory-prey dynamics and community structure and function in such systems [9-10, 22]. However, the results of this study suggest that feeding performance should be taken into account with other biological impacts when predicting the effects of ocean acidification on community and ecosystem structure and function.

Recent evidence suggests that the negative effects of acidification on marine calcification may be offset by increasing food availability [12-13]. While this may be true for some species and life history stages, it appears that the degree of acidification [13] and long-term impacts to feeding ability could influence whether or not certain organisms will be able to use food adequate food to compensate for the negative effects of acidification. The results of this study suggest that arthropods and adult calcifiers may be successful at compensating for acidification effects where food availability is not limited. However, the ability of early life-stage (e.g., larval and juvenile) molluscs and echinoderms to use food as a refuge from such effects could be limited in a more acidified ocean, although more work is needed to confirm or refute this hypothesis.

In this study, I did not test for effects on other biological processes involved in the acquisition of energy (i.e., energy budgets). For example, digestion and stomach size can play a key role in energy acquisition and can be impacted by acidification [23-24]. Indeed studies have reported on the effects of ocean acidification on organismal energy budgets, but are limited [14-17]. As such, more studies should investigate the impacts of ocean acidification on organismal energy budgets under the combined effects of food supply and acidification in order to accurately predict the full impacts of oceanic acidification on marine calcifiers.

While this study suggests that ocean acidification is likely to impact the feeding performance of marine calcifiers, it is certain that other ocean climate stressors (e.g., warming, hypoxia, eutrophication, etc.) will also influence calcifier feeding responses. The meta-analysis of [25] suggested complex effects of acidification and warming on marine organisms, with the combination of these stressors imposing negative (calcification, reproduction, survival), positive (photosynthesis), and neutral effects (growth). Furthermore, the effects of acidification and warming in combination appear taxonomically-dependent for some, but not all, response variables, with acidification and warming imposing both synergistic and alleviatory effects depending on the taxa and response variable examined [25]. From the papers consolidated in this study, only one study focused on the effects of ocean acidification and hypoxia on calcifier feeding performance [17] while 8 assessed the independent and combinatory effects of ocean acidification and warming on calcifier feeding. Consequently, much more work is needed to better understand the effects of ocean acidification and warming in combination on marine calcifier feeding.

Ultimately, it appears that ocean acidification can influence the feeding performance of marine calcifiers, although such responses are taxon-dependent. More studies should now focus on effects in other marine calcifiers (e.g. corals) and continue to investigate the effects of acidification and warming in combination. Studies should also work toward a more holistic understanding of acidification and warming effects on energy budgets and how such effects fit into models of community and ecosystem effects of oceanic climate change.

## Data accessibility

Supporting data can be found in the supplementary material accompanying this article. Raw data are available upon request.

## Acknowledgements

The author wishes to thank Ms. Victoria Neville and Mr. Brady Quinn for feedback on an earlier version of this manuscript. This research was supported by UNB and NBIF graduate scholarships to the author.

## Funding

This study was funded by UNB and NBIF graduate scholarships to JCC.

## Conflict of interest

None.

